# Wheat bZIPC1 interacts with FT2 and contributes to the regulation of spikelet number per spike

**DOI:** 10.1101/2023.08.01.551548

**Authors:** Priscilla Glenn, Daniel P. Woods, Junli Zhang, Gilad Gabay, Natalie Odle, Jorge Dubcovsky

**Affiliations:** Department of Plant Sciences, University of California, Davis, CA 95616, USA; Howard Hughes Medical Institute, Chevy Chase, MD 20815, USA

**Keywords:** Spike development, spikelet number, wheat, *FLOWERING LOCUS T2* (FT2), bZIP, cereals

## Abstract

Loss-of-function mutations and natural variation in the gene *FLOWERING LOCUS T2* (*FT2*) in wheat have previously been shown to affect spikelet number per spike (SNS). However, while other FT-like wheat proteins interact with bZIP-containing transcription factors from the A-group, FT2 does not interact with any of them. In this study, we used a yeast-two-hybrid screen with FT2 as bait and identified a grass-specific bZIP-containing transcription factor from the C-group, designated here as bZIPC1. Within the C-group, we identified four clades including wheat proteins that show Y2H interactions with different sets of FT*-*like and CEN*-*like encoded proteins. *bZIPC1* and *FT2* expression partially overlap in the developing spike, including the inflorescence meristem. Combined loss-of-function mutations in *bZIPC-A1* and *bZIPC-B1* (*bzipc1*) in tetraploid wheat resulted in a drastic reduction in SNS with a limited effect on heading date. Analysis of natural variation in the *bZIPC-B1* (TraesCS5B02G444100) region revealed three major haplotypes (H1-H3), with the H1 haplotype showing significantly higher SNS, grain number per spike and grain weight per spike than both the H2 and H3 haplotypes. The favorable effect of the H1 haplotype was also supported by its increased frequency from the ancestral cultivated tetraploids to the modern durum and common wheat varieties. We developed markers for the two non-synonymous SNPs that differentiate the *bZIPC-B1b* allele in the H1 haplotype from the ancestral *bZIPC-B1a* allele present in all other haplotypes. These diagnostic markers are useful tools to accelerate the deployment of the favorable *bZIPC-B1b* allele in pasta and bread wheat breeding programs.

**Key Message:** The wheat transcription factor bZIPC1 interacts with FT2 and affects spikelet and grain number per spike. We identified a natural allele with positive effects on these two economically important traits.

## INTRODUCTION

Wheat contributes to almost 20% of the world caloric intake, with approximately 770 million tons produced annually across the globe (fao.org, marketing year of 2020/21). Current rates of yield increase (0.9%) are insufficient to match the projected growth of the human population (Ray et al. 2013), which has generated a renewed interest in understanding and improving this trait. However, yield is difficult to study because it is determined by multiple genes with complex epistatic interactions and interactions with the environment (Reynolds et al. 2012). To facilitate its genetic dissection, total grain yield can be divided into several yield components, including number of spikes per surface unit, grains per spike (GNS) and thousand kernel weight (TKW). The number of grains per spike can be further divided into spikelet number per spike (SNS) and grains per spikelet (also known as fertility).

Of the different yield components, SNS has the highest heritability (Zhang et al. 2018) which facilitates its genetic characterization. The number of spikelets per spike is determined when the inflorescence meristem (IM) stops producing lateral meristems and transitions into a terminal spikelet. This transition occurs early after the initiation of the reproductive phase, limiting the influence of the environment later in the growing season and resulting in a higher heritability than other grain yield components that are affected by the environment throughout the growing season. Increases in SNS have been associated with increases in grain yield in productive genotypes grown in optimum environments (Boden et al. 2015; Dobrovolskaya et al. 2014; Hai et al. 2008; Kuzay et al. 2019; Wolde et al. 2019). However, in suboptimal environments or in source-limited genotypes the increases in SNS are no longer associated with yield increases due to reductions in fertility and/or kernel weight (Kuzay et al. 2019).

In wheat, SNS has been shown to be influenced by flowering time, with genotypes flowering earlier usually having lower SNS than those flowering later (Shaw et al. 2013). Flowering time in wheat is mainly regulated by the vernalization (long exposures to low temperatures providing the competency to flower) and photoperiod pathways (requirement for long days), which converge on the regulation of *FT1* (Distelfeld et al. 2009), the wheat homolog of *Arabidopsis thaliana* (Arabidopsis) *FLOWRING LOCUS T* (*FT*) and *Oryza sativa* (rice) *HEADING DATE 3* (*Hd3*) (Corbesier et al. 2007; Distelfeld et al. 2009; Tamaki et al. 2007). The FT-encoded protein, also referred to as florigen, has been shown to be a mobile signal transported through the plant phloem from the leaves to the shoot apical meristem (SAM) in both Arabidopsis and rice (Corbesier et al. 2007; Tamaki et al. 2007).

In wheat, *FT1* is also a strong promoter of flowering that is expressed in the leaves and not in the SAM, so it is also assumed to be a mobile signal (Lv et al. 2014; Yan et al. 2006). In the absence of *FT1* expression (e.g. under short days in photoperiod sensitive wheats), the SAM transitions to the reproductive phase but spike development is arrested, and spikes fail to emerge from the leaf sheaths or head extremely late (Shaw et al. 2020). In rice, it has been shown that Hd3 activates the expression of MADS-box meristem identity genes by forming complexes with 14-3-3 and FD-like proteins (Florigen Activation Complex), which bind to the MADS-box gene promoters (Taoka et al. 2011). In wheat, FT1 has also been shown to interact with the bZIP transcription factor FDL1 and a variety of 14-3-3 proteins which are thought to act as a bridge facilitating the interaction between FT1 and FD-like proteins (Li et al. 2015; Li and Dubcovsky, 2008). Once FT1 arrives at the SAM, the floral activation complex plays a critical role in accelerating the transition from the vegetative to the reproductive phase by activating floral homeotic genes such as *VERNALIZATION1* (*VRN1*) (Li et al. 2015).

The expansion of the *FT-*like gene family in monocot and eudicot species has led to the diversification of *FT* functions (Ballerini and Kramer, 2011; Chardon and Damerval, 2005; Higgins et al. 2010; Pin et al. 2010; Woods et al. 2019). In wheat, twelve different *FT-*like genes have been identified (Halliwell et al. 2016; Lv et al. 2014) and evidence of sub-functionalization has been found. For example, *FT3* and *FT5* have a more critical role in the induction of flowering under SD than other LD expressed *FT*-like genes (Faure et al. 2007; Kikuchi et al. 2009; Lv et al. 2014; Zikhali et al. 2017). In addition, *FT2* was shown to have a more predominant influence on SNS than on heading time (Glenn et al. 2022; Shaw et al. 2019).

The functional differences in *FT2* are particularly interesting, because this gene is the closest paralog of *FT1* (78% identical at the protein level) and the only *FT-*like gene in wheat that has been shown to be transcribed directly in the SAM and developing spike (Shaw et al. 2019). Loss-of-function mutations in both homoeologs of *FT2* in tetraploid wheat (henceforth *ft2*) result in small differences in heading time but significant increases in SNS. However, the *ft2* mutant also shows reduced fertility precluding its utilization in breeding applications (Shaw et al. 2019). Fortunately, a natural variant of *FT-A2*, resulting in an Aspartic Acid (D) change to Alanine (A) at position 10 (henceforth, D10A), has been recently associated with increases in SNS, GNS and grain weight per spike (GWS) indicating no negative impacts on fertility (Glenn et al. 2022). The beneficial A10 allele was almost absent in tetraploid wheat but was present in more than half of the analyzed common wheat varieties suggesting that this allele has been favored by selection for improved grain yield in common wheat (Glenn et al. 2022).

The functional differentiation of *FT1* and *FT2* was paralleled by differences in their protein-protein interactions. Whereas FT1 was able to interact in yeast-two-hybrid (Y2H) assays with five of the six tested 14-3-3 proteins, FT2 showed no interactions with any of them (Li et al. 2015). The two proteins also differed in their interactions with FDL proteins. Specifically, FT1 showed positive Y2H interactions with FDL2 and FDL6, in contrast FT2 showed a positive interaction with FDL13 (Li and Dubcovsky, 2008). However, later it was found that FDL13 was an alternative splice form of FDL15 with a retained intron and a premature stop codon (Li et al. 2015), and the complete FDL15 did not interact with FT2. In summary, no protein interactors of FT2 were identified before this study.

Here, we explored the proteins FT2 can interact with by performing a Y2H screen and identified a bZIP-containing transcription factor from the grass-specific C-group that we designated as *bZIPC1*. We show that the proteins encoded by *bZIPC1* and three other members of the C-group can interact with FT*-* and CEN-like wheat proteins, a function that was previously shown only for bZIP members of the A-group (FDL2, FDL6 and FDL15) (Li et al. 2015). Functional characterization of *bZIPC1* mutants revealed that this gene impacts SNS but has limited effect on heading time. Finally, we characterized natural variation in this gene and identified variants associated with differences in SNS that may be of value for wheat breeding applications.

## MATERIALS AND METHODS

### Plant materials and growth conditions

We identified the *bZIPC1* orthologs in the durum wheat cultivar Kronos using the sequences from Chinese Spring *bZIPC-A1* (TraesCS5A02G440400) and *bZIPC-B1* (TraesCS5B02g444100) and identified loss-of-function mutations in the sequenced Kronos mutant population (Krasileva et al. 2017). We then combined mutations in the A and B homoeologs from each gene to generate three loss-of-function lines designated as *bzipc1-1*, *bzipc1-2* and *bzipc1-3*.

*bZIPC1* mutant lines were initially grown in growth chambers at 16 hour long-day with temperatures oscillating between 22 and 18 °C during the day and night, respectively. After heading and phenotyping, plants were moved to a greenhouse for drying and seed increases.

Seed increases were grown in long-day greenhouses given supplemental lighting during the winter.

### Phylogenetic and statistical analyses

Phylogenetic analyses of bZIP C-group genes were performed using *BdbZIPC1*, *AtbZIP9*, *AtbZIP10*, *AtbZIP25* and *AtbZIP63*, as seed sequences for BLAST searches using Phytozome and NCBI as described previously (Woods et al. 2011). Amino acid sequences were aligned using MUSCLE (Edgar, 2004) before a manual alignment of amino acid sequences in Mesquite (Maddison and Maddison, 2007; Fig. S1). Unalignable regions were pruned from the analysis to minimize noise in the phylogenetic signal, and only 83 contiguous amino acids including the basic region and the leucine zipper domain were used in the final analysis. The Neighbor-Joining method (Saitou and Nei,1987) was used to infer the evolutionary history of the bZIP C-group across flowering plant diversification using five Arabidopsis bZIP proteins from the S-group as outgroups. The percentage of replicate trees in which the associated taxa clustered together in the bootstrap test (1000 replicates) are shown next to the branches (Felsenstein 1985). The evolutionary distances were computed using the Poisson correction method and are presented in number of amino acid substitutions per site. Evolutionary analyses were done using SeaView version 4 (Gouy et al. 2010) and further annotated in Adobe Illustrator 2023.

Analysis of Variance was conducted with the “Anova” function in R package “car” (Fox and Weisberg, 2020) with type 3 sum of squares and LS Means to accommodate unbalanced designs.

### Yeast two-hybrid (Y2H) assays

The full-length coding region of *FT2* (TraesCS3A02G143100) was cloned from Chinese Spring as described in Li et al. (2015). The *FT2* coding region was recombined into pDONRzeo using Life Technologies BP Clonase following the manufacturer’s protocol. *FT2* in pDONRzeo was subsequently recombined into the pDEST32 and pDEST22 yeast destination vectors using Life Technologies LR Clonase II following the manufacturer’s protocol. Clones were verified by sequencing at each cloning step to ensure sequence integrity. All direct assays and library screens were performed using the MaV203 yeast strain as described in the ProQuest manual (Invitrogen). Before screening the cDNA libraries, we confirmed that FT2 was not auto-activated when used as prey or bait by testing it against the empty vector.

We screened a cDNA “photoperiod” library previously developed in *B. distachyon* (Cao et al. 2011). The photoperiod library was generated from shoots of two-week-old plants collected over 24 h grown under long (20 h light) and short day (8 h light) photoperiods (Cao et al. 2011). For the screen, transformants were selected on plates with synthetic minimal media lacking leucine (L) and tryptophan (W) and were replica plated on synthetic minimal media lacking L, W and uracil (U) to identify putative FT2 interactors. Yeast colony polymerase chain reaction (PCR) was done using the Phire polymerase following manufacturer’s instructions (Thermo Fisher).

To confirm the Y2H screen results, we cloned the full-length coding sequence from Bradi1g05480. The complete *bZIPC1* coding region was amplified by PCR and cloned into the pJET vector (ThermoFisher), and from there amplified using bZIPC1 BP primers (Table S1) and recombined into pDONRzeo using Life Technologies BP clonase. The pDONRzeo vector containing the desired *bZIPC1* coding region was recombined into pDEST22 and pDEST32 using Life Technologies LR Clonase II. Clones were verified by sequencing at each cloning step. We also cloned the wheat *bZIPC1* paralogs *bZIPC3* and *bZIPC4*, and genes *CEN4*, *CEN5*, *FT2*, *FT3* and *FT5* using the same cloning strategy described for *bZIPC1* (Table S1), whereas *bZIPC2* was synthesized and cloned. *bZIPC1* and all genes used in the directed Y2H assays were confirmed to not auto-activate when either used as prey or bait. *FT1* and *CEN2* were previously cloned (Li et al. 2015) and transferred into pDEST22 and pDEST32.

### Spatial Transcriptomics

Between the two homoeologs present in tetraploid wheat for each gene, we selected the one expressed at higher levels in the Kronos transcriptome of the developing spike (VanGessel et al. 2022): *bZIPC-A1* (TraesCS5A02G440400), *bZIPC-B3* (TraesCS5B02G059200), *bZIPC-A4* (TraesCS6A02G154600) and *FT-A2* (TraesCS3A02G143100). *bZIPC2* (TraesCS1A02G329900 and TraesCS1B02G343500) was excluded from this study because it is expressed primarily during grain development (Choulet et al. 2014) and was not detected in the Kronos spike transcriptome study (VanGessel et al. 2022). We also provided Resolve Biosciences the sequences of the homoeologs of the selected genes, and they excluded them in their quality control for specificity performed against all the coding sequences of the wheat genome (Ref Seq v1.1). Therefore, although probes were designed based on the sequence of the highest expressing homoeolog, they are not genome-specific and can detect both homoeologs for each gene.

Probes for *bZIPC1*, *bZIPC3*, and *bZIPC4* passed the quality and specificity thresholds but, initially, it was not possible to develop specific probes for *FT2* because of its close similarity with *FT1*. We then excluded the sequences of *FT-A1* and *FT-B1* from the reference genome used for the specificity check, and Resolve Biosciences was able to design probes to differentiate *FT2* and *FT1* from other wheat genes. Although these probes can potentially detect *FT1*, the lack of expression of *FT1* in the Kronos developing spike (VanGessel et al. 2022) ensures that the signals presented for this tissue are specific for *FT2*.

### Haplotype analysis and marker development

We performed a haplotype analysis using the exome capture data from 55 tetraploid and hexaploid wheat accessions from the genotyping project “2017_WheatCAP_UCD” (https://wheat.triticeaetoolbox.org/downloads/download-vcf.pl) available in the Wheat T3 database (Blake et al. 2016). Using the 242 SNPs detected in a 1.3 Mb region on chromosome 5B flanking *bZIPC-B1* (CS RefSeq v1.1 615,695,210 to 617,038,639), we performed a cluster analysis using R functions ‘dist’ and ‘hclust’ (method = ‘ward.D2’) (R Core Team, 2021)

We used the available exome capture sequence from Berkut and Patwin-515HP (Wheat/T3) to develop KASP markers for the *bZIPC-B1* SNPs. We also created KASP markers for the induced mutant alleles (Table S1).

## RESULTS

### Identification of FT2 interactors by Y2H screening

To identify proteins that interact with FT2, we performed Y2H screens using the wheat Chinese Spring FT-A2 protein (TraesCS3A02G143100) as bait and a *B. distachyo*n photoperiod cDNA library developed from harvesting whole plants across a diurnal cycle in long and short photoperiods (Cao et al. 2011) as prey. In the three screens performed against this library (transformation efficiencies of 4.5x10^5^, 4.6X10^6^ and 4.1x10^6^), we detected 121 positive colonies corresponding to 26 different genes (Table S2). Among them, the gene represented by the largest number of colonies (67) was *Bradi1g35230* (annotated as a kinectin-related protein with motor activity). The second most abundant gene was *Bradi1g05480*, which was detected in 29 colonies. This gene was annotated as an uncharacterized bZIP transcription factor. We decided to prioritize this bZIP for functional analyses because it was the only interactor detected in all three Y2H screens and also because of the known protein-protein interactions between FT1 and other bZIP proteins (Li et al. 2015). The interactors detected in the Y2H screens and their functional annotation are presented in Table S2.

### Phylogenetic analyses of *bZIPC*-like genes

The large family of bZIP domain transcription factors includes 13 subfamilies (A-L and S) (Guedes Corrêa et al. 2008; Peviani et al. 2016). Reciprocal BLAST searches between Bradi1g05480 and Arabidopsis did not reveal a clear one-to-one orthologous relationship but were sufficient to place Bradi1g05480 within the C-group (Fig. 1), which includes Arabidopsis genes *bZIP9* (At5g24800), *bZIP10* (At4g02640), *bZIP25* (At3g54620), and *bZIP63* (At5g28770). To reflect Bradi1g05480’s phylogenetic relationship within the C-group of this large and diverse family of bZIP transcription factors, we designated this gene as *bZIPC1* (Table 1).

**Fig. 1.**
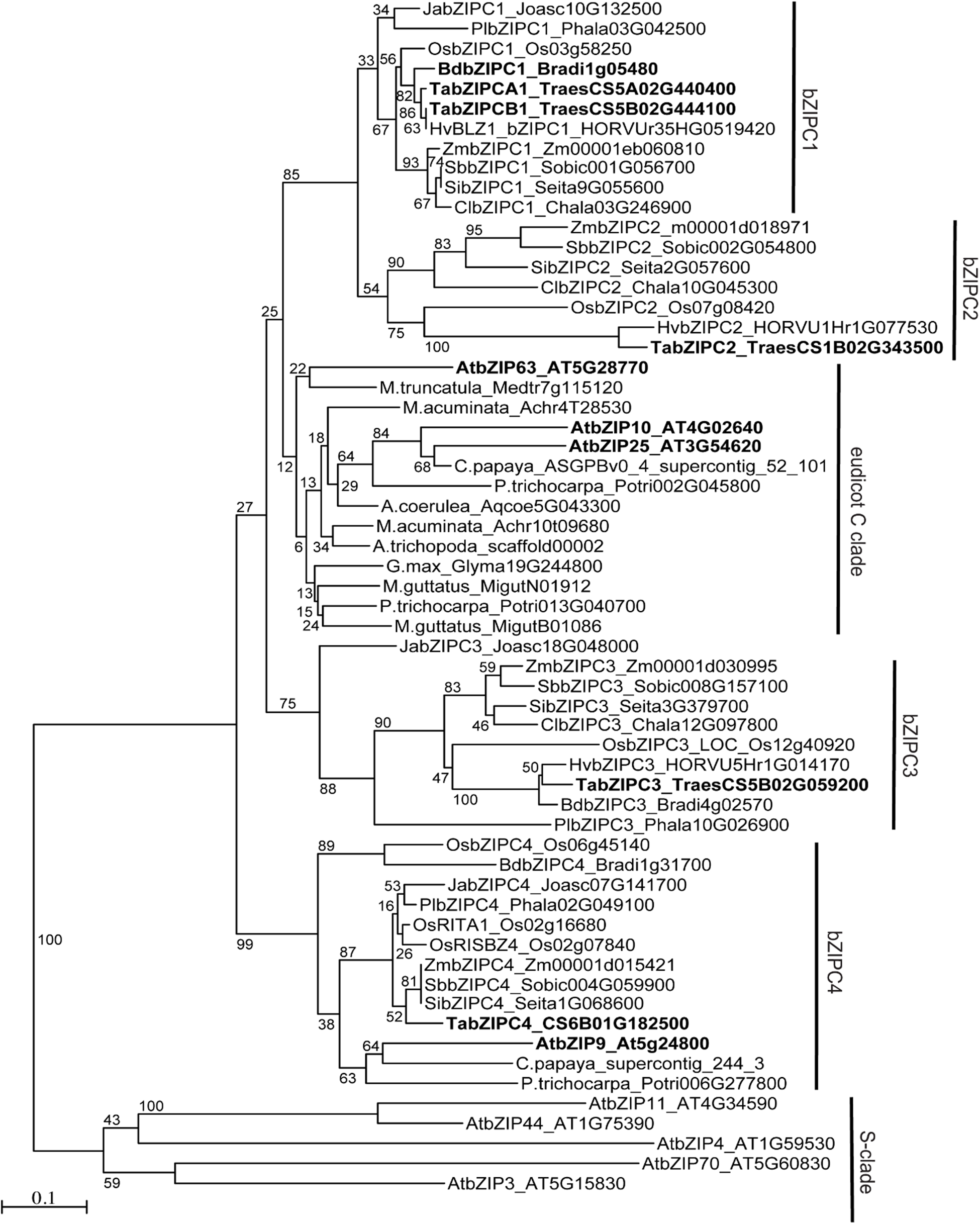
Phylogenetic analysis of 55 C-group and 5 S-group bZIP proteins. The tree was constructed using the Neighbor Joining method using an alignment of 83 amino acids (Fig. S1). Bootstrap values are indicated at the branching points. Bar indicates substitutions per site. Genes discussed in the manuscript are labeled in bold font. At=*Arabidopsis thaliana*, Os=*Oryza sativa*, Zm=*Zea mays*, Cl=*Chasmanthium laxum*, Sb=*Sorghum bicolor*, Si=*Setaria italica*, Hv=*Hordeum vulgare*, Ta=*Triticum aestivum*, Bd= *Brachypodium distachyon*, Pl=*Pharus latifolius*, Ja=*Joinvillea ascendens*.

**Table 1.**
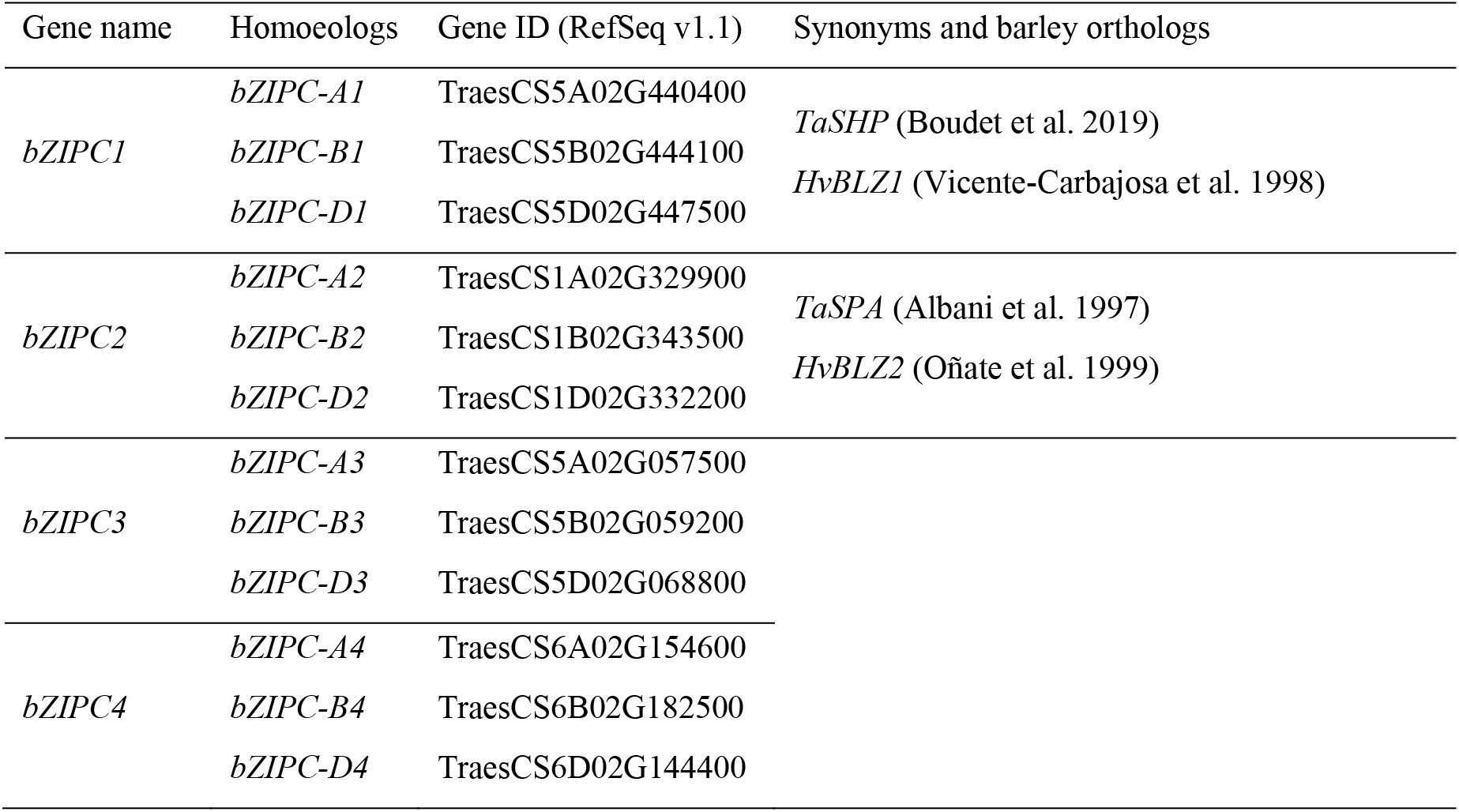
Proposed nomenclature for the *bZIP* wheat genes from the C-group.

To understand the evolutionary history within the bZIP C-group and to identify the closest proteins to Bradi1g05480 in other species, we selected 55 bZIP proteins from the C-group spanning flowering plant diversification and five Arabidopsis bZIP proteins from the close S-group as an outgroup (Fig. 1). Relationships among these C- and S-class bZIP proteins were estimated using Neighbor-Joining phylogenetic methods on an alignment of 83 contiguous amino acids including the conserved basic region and leucine zipper (Fig. S1).

The bZIPC1 clade included grass species and the near relative *Joinvillea* (Fig. 1). The closest clade to bZIPC1, which we designate here as bZIPC2, also included grass species from the major grass subfamilies (Fig. 1). These results suggest that *bZIPC1* and *bZIPC2* are likely the result of a grass specific duplication, which is consistent with the known whole genome duplication event at the base of the grasses (Paterson et al., 2004). These two clades, together with the eudicot clade including the duplicated Arabidopsis bZIP10, bZIP25 and bZIP63 proteins, correspond to the possible orthologous group C1 in Guedes Corrêa et al. (2008).

Two additional clades within the bZIP-C group, designated here as *bZIPC3* and *bZIPC4*, were more distantly related to the bZIPC1/bZIPC2 cluster (Fig. 1). The Arabidopsis C-group protein bZIP9 was included in the same clade as bZIPC4 from grasses, forming a group similar to the proposed orthologous group C3 in Guedes Corrêa et al. (2008). This result suggests that some of the diversification of the bZIP C-group occurred before the monocot - eudicot divide.

To generate a nomenclature for the wheat bZIP transcription factors that reflects their evolutionary relationships, we propose to incorporate the group name to the bZIP gene name and designate the wheat *bZIP* genes from the C-group as *bZIPC1*, *bZIPC2*, *bZIPC3*, and *bZIPC4* (Table 1).

### Interactions between bZIPC-like and FT-like proteins

The *B. distachyon* clones identified in the Y2H library were not full-length genes based on BLASTN and BLASTP comparisons. To validate the interaction with full-length bZIPC1, we conducted direct Y2H assays using the full-length sequence of wheat bZIPC1 protein (TraesCS5A02G440400, 392 amino acids). Since there are two different natural variants in FT-A2 known to impact SNS (D10 and A10) (Glenn et al. 2022), we tested both protein variants in the direct Y2H assays. We found that bZIPC1 can interact with both FT-A2 protein variants in yeast when used as either bait or prey, confirming the initial Y2H screen results (Table 2 and Fig. 2).

**Fig. 2.**
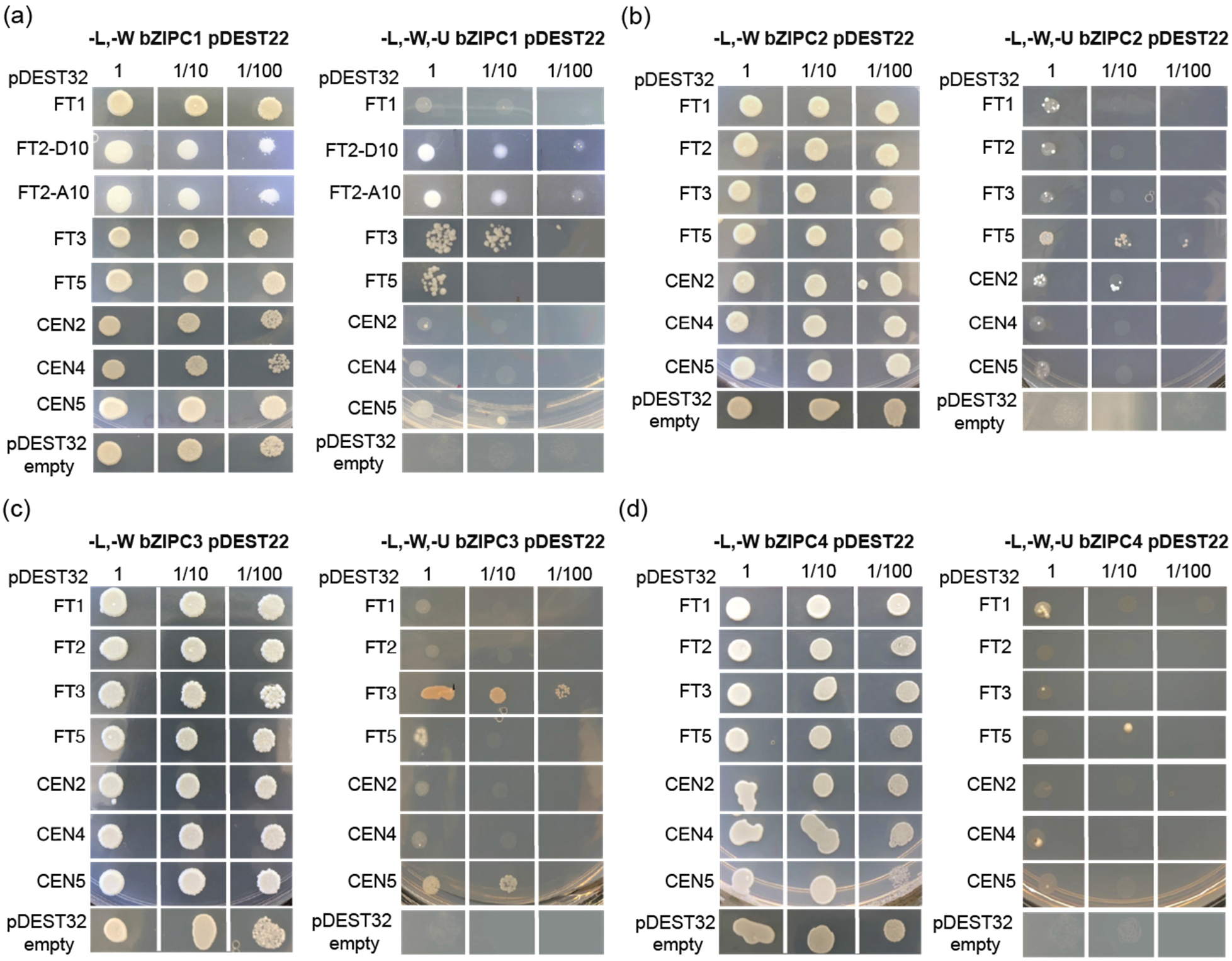
Interactions between bZIPC-like and FT/CEN-like proteins. Yeast two-hybrid assays between (a) bZIPC1 (b) bZIPC2, (c) bZIPC3, (d) bZIPC4 and FT1, FT2-D10, FT2-A10 (bZIPC1 only), FT3, FT5, CEN2, CEN4, CEN5 and empty pDEST32 (auto-activation test). Left panel: SD medium lacking Leucine and Tryptophan (-L-W) to select for yeast transformants containing both bait and prey. Right panel: interaction on -L-W-U medium (no L, W, and Uracil). Dilution factors = 1, 1:10 and 1:100.

**Table 2.** Yeast-two-hybrid interactions between wheat bZIP proteins from the C-group and FT-like and CEN proteins.

| Bait pDEST32 | Prey pDEST 22 |  |  |  |
| --- | --- | --- | --- | --- |
|  | bZIPC1 | bZIPC2 | bZIPC3 | bZIPC4 |
| FT1 | NO | WEAK | NO | WEAK |
| FT2-D10 | YES | WEAK | NO | NO |
| FT2-A10 | YES | NA | NA | NA |
| FT3 | YES | WEAK | YES | NO |
| FT5 | WEAK | YES | WEAK | NO |
| CEN2 | NO | YES | NO | NO |
| CEN4 | NO | WEAK | NO | WEAK |
| CEN5 | WEAK | NO | YES | NO |

To explore if bZIPC1 can interact with other FT-like proteins, we performed pairwise Y2H assays between bZIPC1 and FT-A1 (TraesCS7A02G115400), FT-B3 (TraesCS1B02G351100), FT-B5 (TraesCS4B02G379100), CEN-B2 (TraesCS2B02G310700), CEN-A4 (TraesCS4A02G409200) and CEN-A5 (TraesCS5A02G128600). We found that in addition to FT2, bZIPC1 can interact with FT3 and weakly with FT5 (Table 2 and Fig. 2).

We also tested if the bZIPC1 paralogs were to interact with the same set of FT-like proteins in a set of Y2H pairwise tests (Fig. 2). bZIPC2 interacted strongly with FT5 and CEN2 and weakly with the other genes, excluding CEN5, which showed no detectable interaction (Fig. 2b). bZIPC3 only interacted with FT3 and CEN5 and weakly with FT5; whereas the bZIPC4 protein only interacted weakly with FT1 and CEN4 (Table 2 and Fig. 2).

### Expression of *bZIPC1* and *FT2* overlap in the developing spikelet

We took advantage of published RNAseq data in Chinese Spring from five different tissues at three developmental stages (Choulet et al. 2014) to evaluate the expression levels of the four different *bZIPC* genes. In CS, *bZIPC1* transcripts for the three homoeologs were detected across all tissues with the highest expression in the roots (Fig. S3a). The spike tissues, all collected after the initiation of the elongation phase (Zadoks scale Z32, Z39 and Z65 (Zadoks et al. 1974)), also showed high and consistent expression levels of all three *bZIPC1* homoeologs. *bZIPC2* transcripts were detected only in the developing grains (Fig. S3b), whereas *bZIPC3* transcript levels were low in the developing grains, but abundant in other tissues (Fig. S3c). *bZIPC4* transcripts were detected in all tissues, with higher levels in the spike (particularly at the latest stage Z65) and stems (Fig. S3d).

To study the expression of *bZIPC1* earlier in spike development, we used an RNAseq study in tetraploid wheat Kronos including samples collected at the vegetative (W1.0), double ridge (W2.0), glume primordia (W3.0), lemma primordia (W3.25), and floret primordia (W3.5) stages (VanGessel et al. 2022), with W values based on the Waddington spike developmental scale (Waddington et al. 1983). Overall expression was similar between the A and B genomes, with expression levels increasing from W1.0 to W3.0 and remaining constant through W3.5 (Fig. 3a). *FT2* expression levels in the same RNAseq samples were extremely low in W1.0 but then increased significantly in W2.0 to W3.0 before stabilizing (Fig. 3b). Thus, *bZIPC1* and *FT2* are expressed in the same organ at the same developmental stages.

**Fig. 3.**
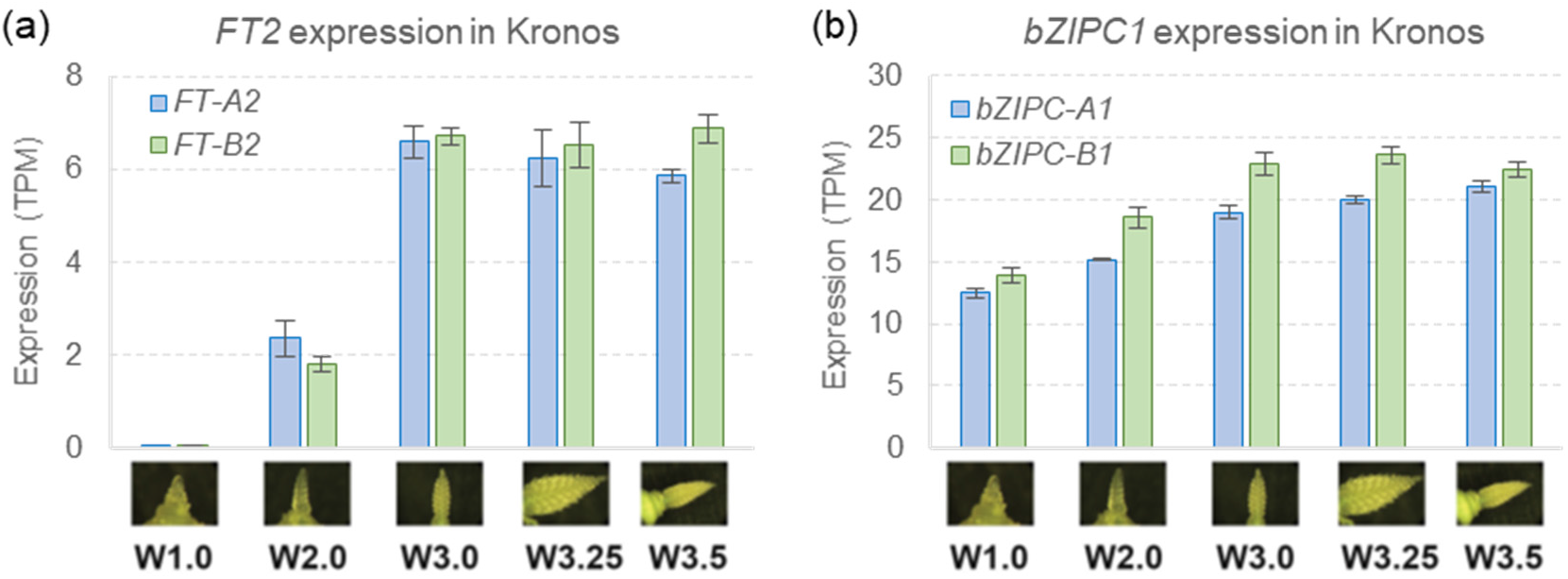
Overlapping expression of *bZIPC1* and *FT2* in the wheat spike. (a) Expression of *bZIPC-A1* (TraesCS5A02G44044) and *bZIPC-B1* (TraesCS5B02G444100) in tetraploid wheat Kronos across the early stages of spike development (n = b). (c) Expression of *FT-A2* and *FT-B2* in tetraploid wheat Kronos across the early stages of spike development (n = 4).

To explore if the *bZIPC1* and *FT2* genes were co-expressed in the same regions of the developing spike, we performed a spatial transcriptomics experiment. Figure 4a shows the distribution of *bZIPC1*, *bZIPC3*, and *bZIPC4* transcripts in a developing spike of tetraploid wheat Kronos at the lemma primordia stage (W3.25). At this stage, *bZIPC1* showed a strong and uniform distribution across the developing spike (Fig. 4a-b). *bZIPC3* also showed a relative strong signal and a similar distribution, except for a reduced presence in the distal region of the spikelet meristems (Fig. 4a). Finally, *bZIPC4* showed a lower signal localized in the central part of the spike (Fig. 4 b-c). The different distribution of the three *bZIPC* genes suggests good specificity of the probes.

**Fig. 4.**
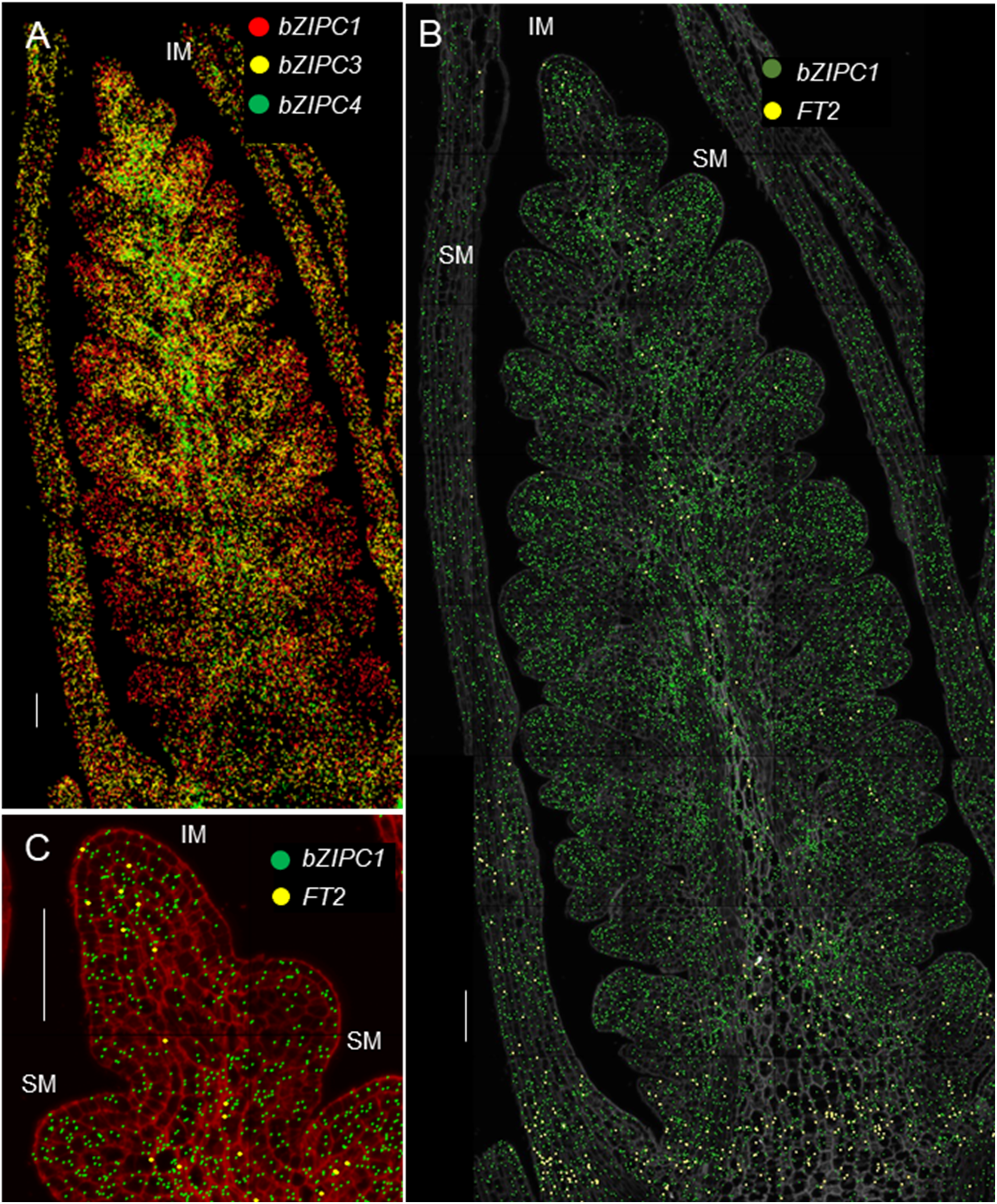
Spatial distribution of *bZIPC* and *FT2* genes in the wheat developing spike. Kronos spike at the lemma primordia stage (W2.25). (a) *bZIPC1* (red), *bZIPC3* (yellow) and *bZIPC4* (green). (b) *bZPC1* (dark green) and *FT2* (yellow). (c) Detail of B showing co-localization of *bZIPC1* and *FT2* within the same cells (cell walls in red). IM = inflorescence meristem, SM = spikelet meristem. Scale = 100 μm

The signal of *FT2* was lower than that of the *bZIPC* genes and was localized mainly at the base and along the central part of the developing spike reaching the IM (Fig. 4b). Therefore, *FT2* overlapped with *bZIPC1* across its distribution, and the two genes were co-localized within the same cells in the IM (Fig. 4c), where the transition to the terminal spikelet is close to occurring (usually at W3.5). These results support the hypothesis that the physical interaction between bZIPC1 and FT2 observed in Y2H can be biologically relevant.

### Loss-of-function mutations in *bZIPC1* are associated with reduced SNS

To determine the function of *bZIPC1*, we combined loss-of-function mutations in the two homoeologs of this gene in tetraploid wheat Kronos. We identified one Kronos mutant line with a loss-of-function mutation in the A genome homoeolog *bZIPC-A1* (TraesCS5A02G440400) and three in the B genome homoeolog *bZIPC-B1* (TraesCS5B02G444100). The mutation in *bZIPC-A1* (CS RefSeq v.1, chromosome 5A position 621,763,602) in the Kronos mutant K3308 results in a premature stop codon at position 97 (Q97*) in the first exon. In *bZIPC-B1*, we identified two independent mutants with identical mutations (K2991 and K3532). This mutation is located in chromosome 5B position 616,654,824 at the acceptor splicing site at the end of intron one (Fig. 5a). Sequencing of the *bZIPC-B1* mRNA products in mutants K2991 and K3532 revealed that the second exon (94 bp) was spliced out, generating a shift in the reading frame and a premature stop codon (Fig. S2). We also identified a truncation mutation in Kronos mutant line K2038 at chromosome 5B position 616,655,991 that results in a premature stop codon (W117*).

**Fig. 5.**
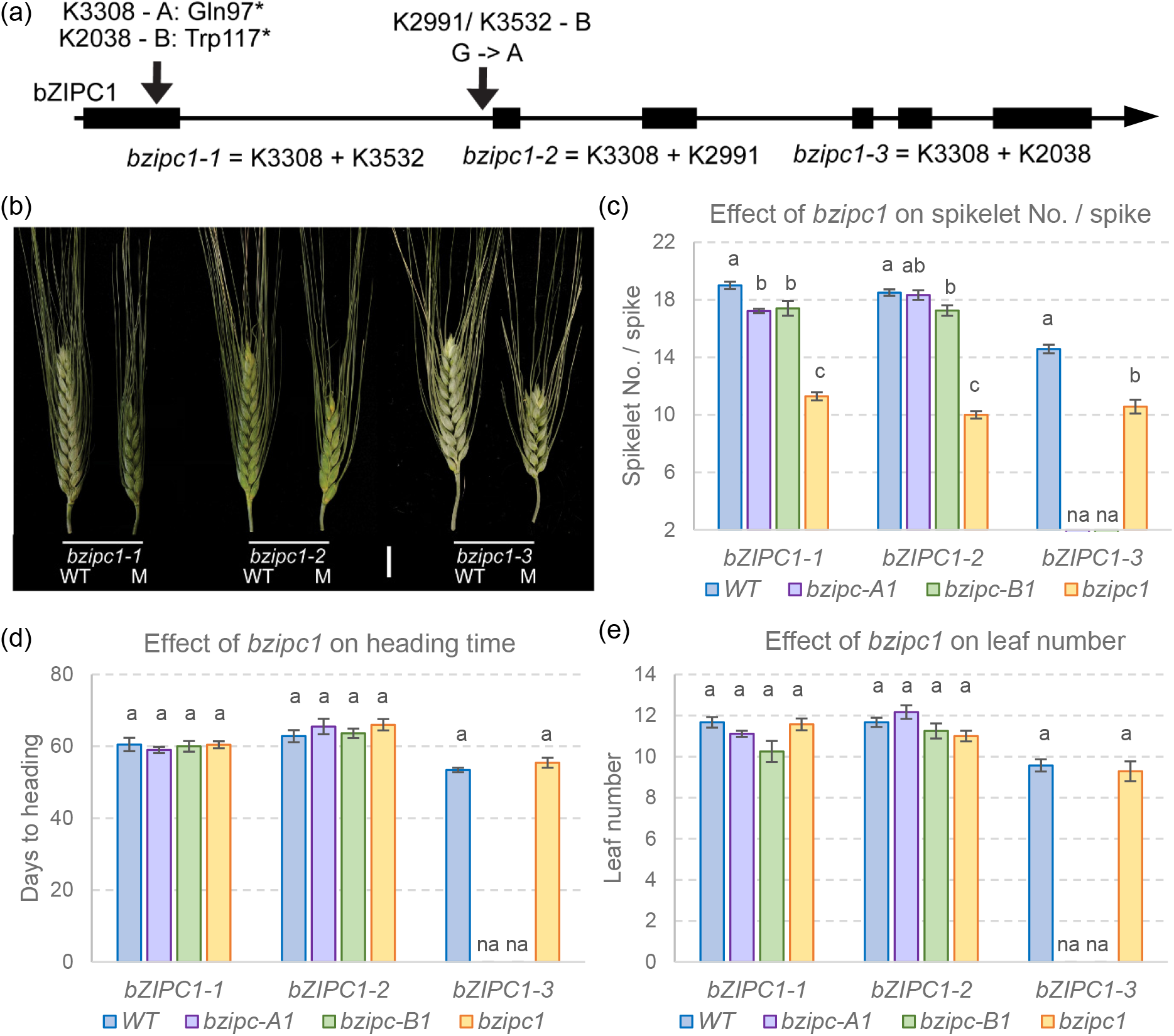
Characterization of F_2_ *bZIPC1* mutants for SNS, and flowering time in a growth chamber. (a) Selected mutations for *bZIPC-A1* and *bZIPC-B1* in Kronos. (b) Spike morphology of WT and mutant sister lines for *bzipc1-1*, *bzipc1-2* and *bzipc1-3*. Scale bar = 2cm (c) Spikelet number (No.) / spike (SNS). (d) Days to heading. (e) Leaf number at heading time. (c-e) Different letters above bars indicate significant differences in Tukey tests (*P* < 0.05). n = 5 to 9.

We designed KASP markers for each of the EMS-induced mutations (Table S1) to trace them during the experiments. We made crosses between the plants carrying the A genome mutation with those carrying the three independent B genome mutants. In the three F_2_ progenies, we selected plants homozygous for the different mutant combinations using the KASP markers (Table S1). We designated the three combined mutants as *bzipc1-1* (K3308 + K3532), *bzipc1-2* (K3308+ K2991), and *bzipc1-3* (K3308 + K2038, Fig. 5a).

All three double mutants showed highly significant reductions in SNS (*P* < 0.001) compared to their wildtype sister lines (average 6.7 spikelets / spike across the three mutants, Table S3), but the individual spikelets showed no obvious differences (Figs. 5b-c). Significant reductions were also observed in a greenhouse experiment using F_2_ plants (average 4.4 spikelets / spike, Table S3) and in a growth chamber experiment using more advanced F_3_ plants (average 6.9 spikelets / spike, Fig. S4, Table S4). These last two experiments included only the *bzipc1-1* and *bzipc1-2* combined mutants and sister wildtype lines due to seed limitations. Among the single mutants, *bzipc-A1-1*, *bzipc-B1-1* and *bzipc-B1-2* showed modest but significant reductions in SNS relative to the wildtype (Fig. 5c).

In the same three experiments, we found no consistent significant differences in days to heading (DTH) or leaf number at heading (LN) between the mutants and their corresponding wildtype sister lines. These results suggest that *bZIPC1* has a limited effect on the timing of the transition between the vegetative and reproductive phase stages and the duration of the elongation phase (Fig. 5d-e).

### Natural variation in *bZIPC1*

The significant differences in SNS observed in the *bzipc1* mutants motivated us to look at natural variation in both *bZIPC1* homoeologs in a set of 6 tetraploid and 49 hexaploid wheat sequenced by exome capture (Fig. 6). We found two non-synonymous SNPs in *bZIPC-B1* (TraesCS5B02G444100) but none in *bZIPC-A1* (TraesCS5A02G440400) or *bZIPC-D1* (TraesCS5D02G447500) in hexaploid wheat, so we focused our haplotype analysis in a 1.3 Mb region on chromosome 5B flanking *bZIPC-B1* (CS RefSeq v1.1 615,695,210 to 617,038,639).

**Fig. 6.**
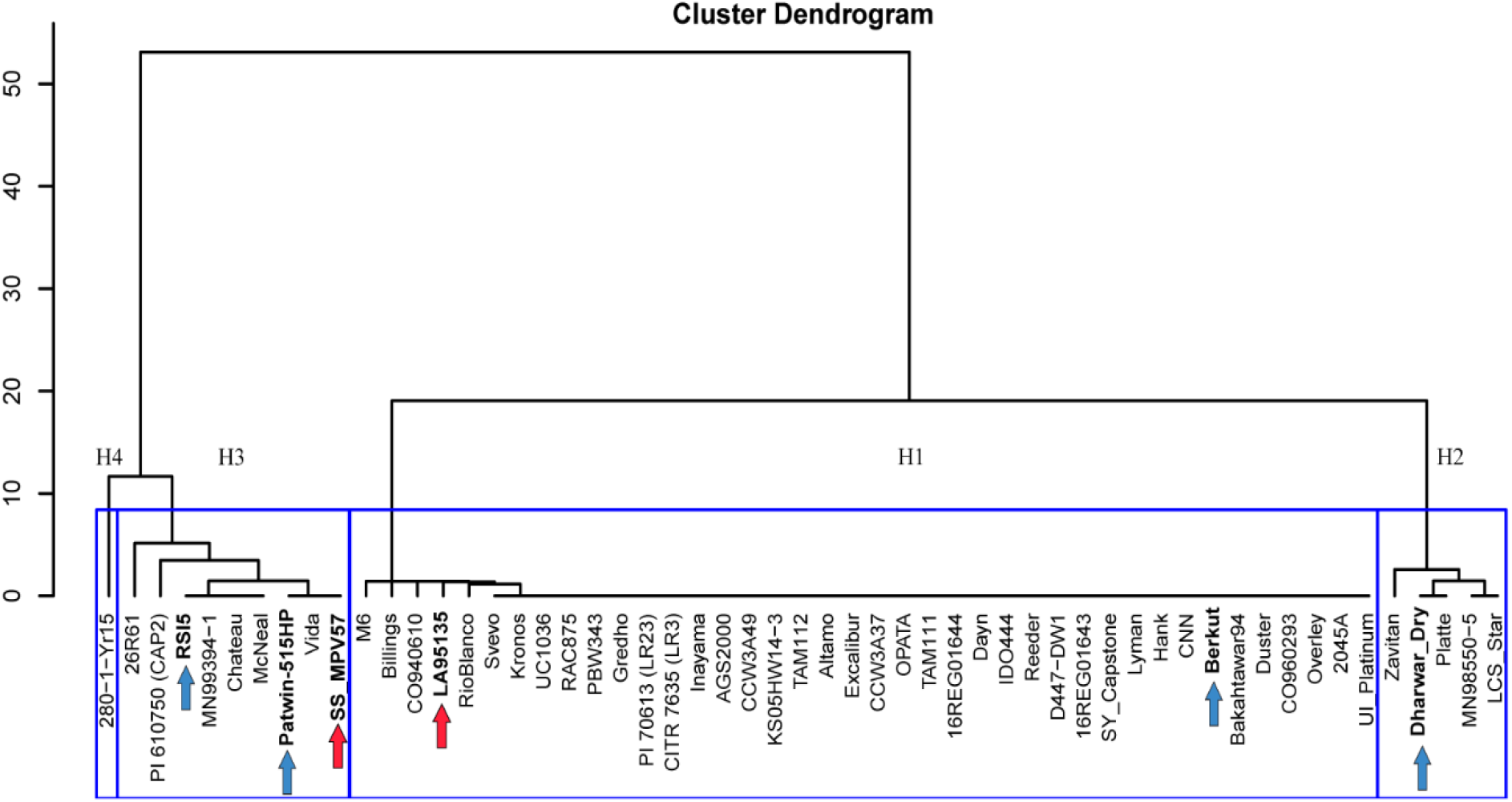
Haplotypes in the *bZIPC1* region. Cluster analysis based on 242 SNPs detected in chromosome 5B (CS RefSeq v1.1 615,695,210 to 617,038,639 bp) in exome capture data extracted from T3/Wheat (https://wheat.triticeaetoolbox.org/). Blue arrows indicate spring lines from haplotypes H2 and H3 used as parental lines in crosses Berkut (H1). Red arrows indicate winter lines used as parental lines in the soft red winter mapping population (DeWitt et al. 2021). Clustering scale calculated from method “ward D2”.

A cluster analysis based on the 242 SNPs detected in this region (Table S5) revealed four haplotypes designated here as H1 to H4 (Fig. 6). The H1 haplotype was the most frequent (72.7%) among these accessions and was more closely related to H2 than to the other two haplotypes. Haplotypes H3 and H4 were related to each other, but since H4 included only a single tetraploid accession (280-1-Yr15) used for the introgression of *Yr15* from *T. turgidum* subsp. *dicoccoides* (Yaniv et al. 2015), we did not characterize it further. We identified three independent historical recombination events among haplotypes within this 1.34 Mb region in varieties 26R61, LA95135, Kronos and Gredho (summarized in Table S5).

We then compared the four haplotypes for polymorphisms within the *bZIPC-B1* coding region and identified one synonymous SNP at position 616,652,860 that differentiated H3 from all other three haplotypes, and two missense SNPs at positions 616,654,272 (N151K) and 616,654,229 (V166M) that differentiate the H1 haplotype (Table S6). The bZIPC-B1 linked amino acids K151 and M166 (KM henceforth) were found only in the H1 haplotypes whereas the amino acid combination N151 and V166 (NV henceforth) were found in the other three haplotypes, *bZIPC-A1*, *bZIPC-D1*, and *Hordeum vulgare*, suggesting that NV is the ancestral state and KM the derived state. In addition, while we observed variation at position 166 (Table S6), the N151 amino acid was conserved in *B. distachyon*, rice, *Setaria*, and *Panicum* supporting the previous hypothesis.

The H1 haplotype was detected in the genomes of tetraploid wheat Svevo (Maccaferri et al. 2019) and Kronos (https://opendata.earlham.ac.uk/opendata/data/Triticum_turgidum/EI/v1/) and PanGenome hexaploid varieties Chinese Spring, Lancer, SY Mattis, Stanley, Jagger, and CDC Landmark (Walkowiak et al. 2020). A comparison of the *bZIPC-B1* promoter region including 1,500 bp upstream of the start codon revealed no polymorphisms among these diverse accessions, suggesting that H1 may be a relatively recent haplotype. A comparison of the *bZIPC-B1* promoter region (1500 bp) between the H1 and H2 haplotypes (ArinaLrFor, Julius, Cadenza, Paragon, and Clair) revealed 9 SNPs and 4 indels, whereas a comparison between the H1 and H3 haplotypes (Norin61, Spelt, Mace, and Rubigus) revealed 19 SNPs and two indels, confirming the closer relationship between the H1 and H2 haplotypes relative to H3.

To explore the variation of the *bZIPC-B1* haplotypes in time, we screened a collection of 63 *T. turgidum* subsp. *dicoccoides*, 77 *T. turgidum* subsp. *dicoccon*, and 303 *T. turgidum* subsp. *durum* with KASP markers for the N151K and V166M SNPs (primers bZIPC1-4229 and bZIPC1-4272, Table S1). We found that 35 % of the *T. turgidum* subsp. *dicoccoides* accessions and 4 % of the *T. turgidum* subsp. *dicoccon* accessions have the derived KM allele (Table 3 and Table S7). For the *T. turgidum* subsp. *durum,* we split the group into old landraces and more modern cultivars and breeding lines. Of the 175 landraces, we found that 37 % had the derived KM allele while of the 128 cultivars and breeding lines, 65 % carried the KM allele (Table 3 and Table S7), indicating an increased frequency of the derived allele in the modern cultivated durum wheats.

**Table 3.**
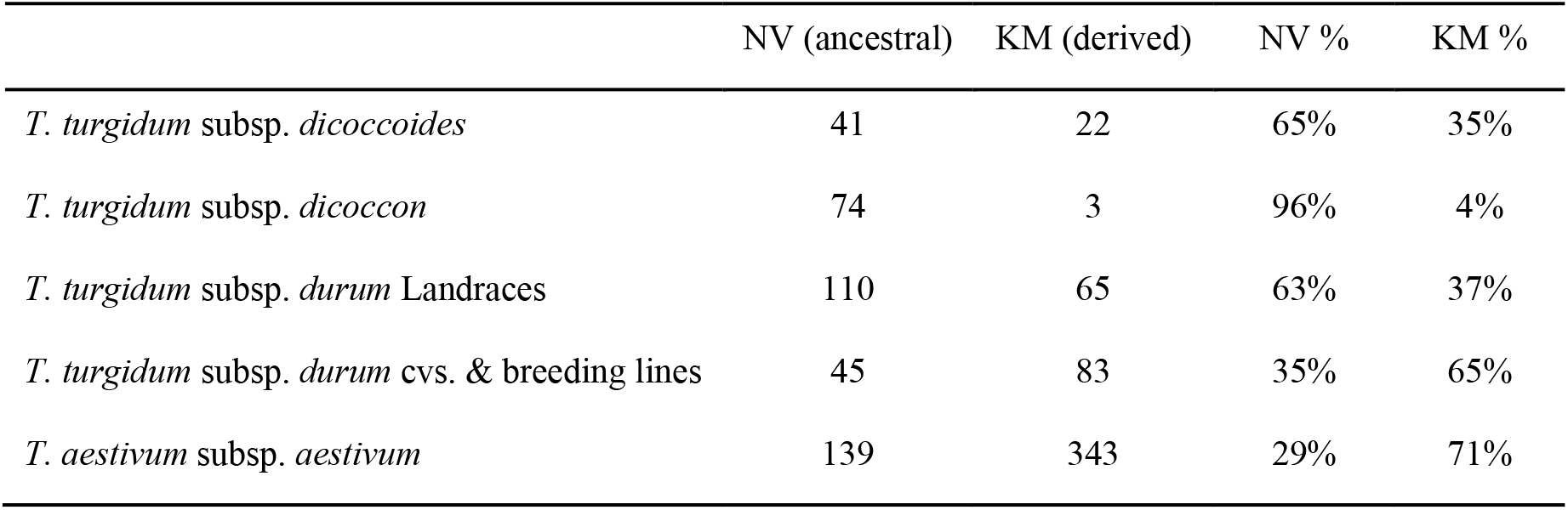
*bZIPC-B1* allele distribution in wild and cultivated tetraploid and hexaploid wheat.

A similar analysis of exome capture data from 482 *T. aestivum* subsp. *aestivum* accessions (He et al. 2019) revealed that 71 % of the accessions have the derived *bZIPC-B1* KM allele and only 29 % the ancestral haplotype (Table 3 and Table S7). Taken together, these results suggest positive selection for the derived KM allele in modern wheat breeding programs. To explore this hypothesis further, we tested the effect of the *bZIPC-B1* KM allele in four segregating populations.

### Effect of *bZIPC1* alleles on grain yield components

The haplotype analysis revealed that the group of varieties with the H1 haplotype included the spring wheat variety Berkut, which is the central parent in the spring wheat Nested Association Mapping (NAM) population (Blake et al. 2019). We previously characterized populations generated from crosses between Berkut (H1) and spring common wheat accessions Patwin-515HP (H3, Table S8), RSI5 (H3, Table S9), and Dharwar Dry (H2, Table S10) for SNS and other grain yield components in multiples locations and under two watering regimes (Zhang et al. 2018).

These populations were previously genotyped with the Illumina 90K SNP chip (Jordan et al. 2015) and among these SNPs, we identified IWB56221 (RefSeq v1.1 5B 616,652,623) as the closest diagnostic marker for the H1 haplotype (H1= T, H2, H3 and H4 = C). Using this marker, we performed *t*-tests between the two alleles for the different traits in the different environments (Table 4).

**Table 4.**
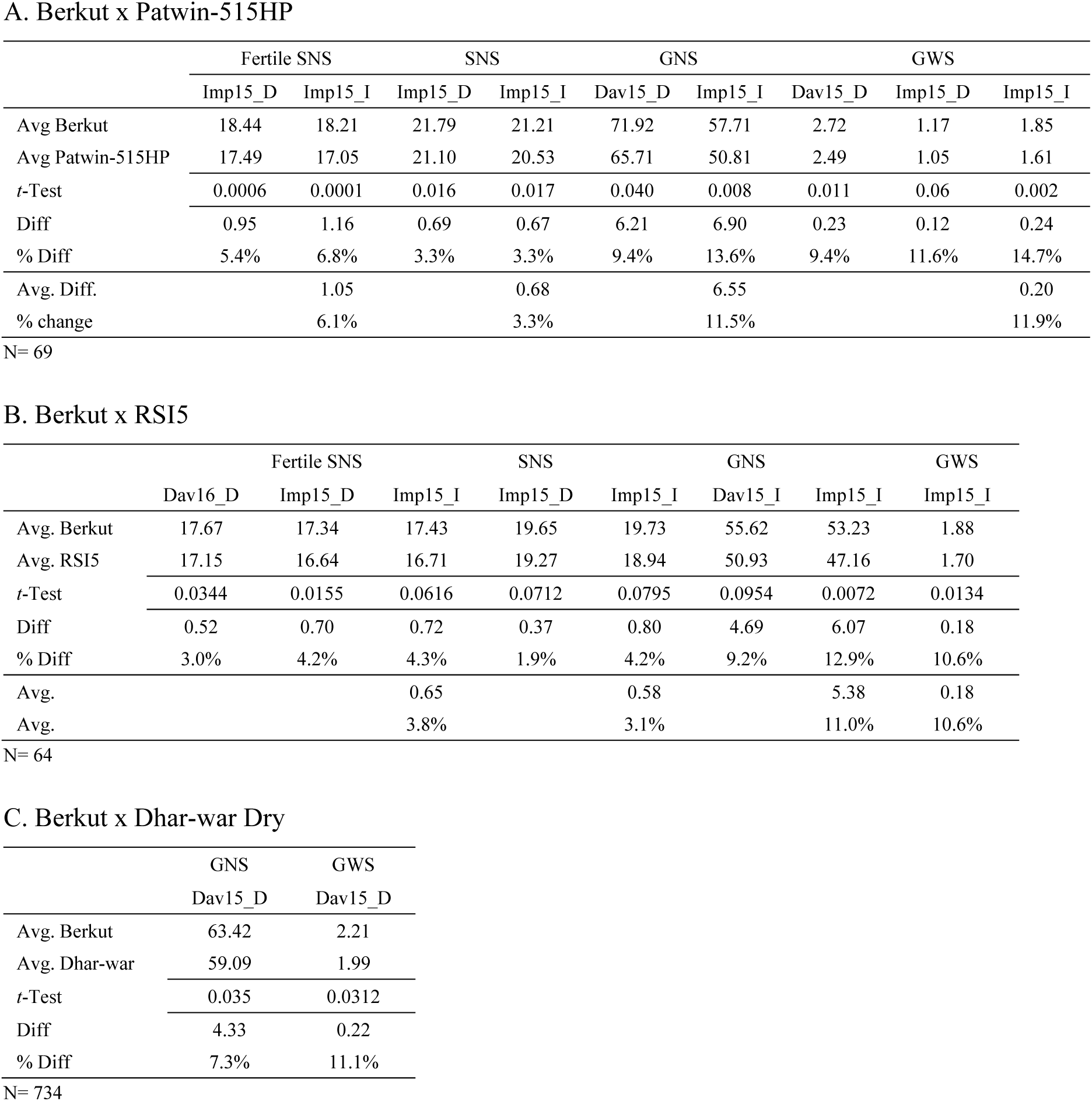
Effect of marker IWB56221 tightly linked to *bZIPC-B1* on total and fertile spikelet number per spike (SNS), grain number per spike (GNS) and grain weight per spike (GWS) in spring wheat NAM populations. Only environments showing significant effects are presented (complete information is available in Supplemental Appendix 2). Imp = Imperial Valley, California, Dav= Davis, Sacramento Valley, California, 15= 2015, 16=2016, I= normal irrigation, and D= without last irrigation. Data is from Zhang et al. 2018.

The comparison between the H1 and H3 haplotypes in the crosses Berkut x Patwin-515HP and Berkut x RSI5 showed that the H1 haplotype (=KM *bZIPC-B1* allele) was associated with significant increases in the number of total and fertile spikelets per spike (3.1 to 6.1%), grain number per spike (11.0 to 11.5% increase), and grain weight per spike (10.6% to 11.9%, Table 5) relative to H3 (NV) in multiple locations. In the cross Berkut x Dharwar Dry, the H1 haplotype was also associated with a significant increase in grain number per spike (7.3%) and grain weight per spike (10.6%, Table 4) relative to the H2 haplotype (NV). Although the number of total and fertile spikelets per spike was not significant in this population, it showed the same trend as in the other two populations, with higher values in H1 (1.1 to 1.7%) relative to H2.

**Table 5.**
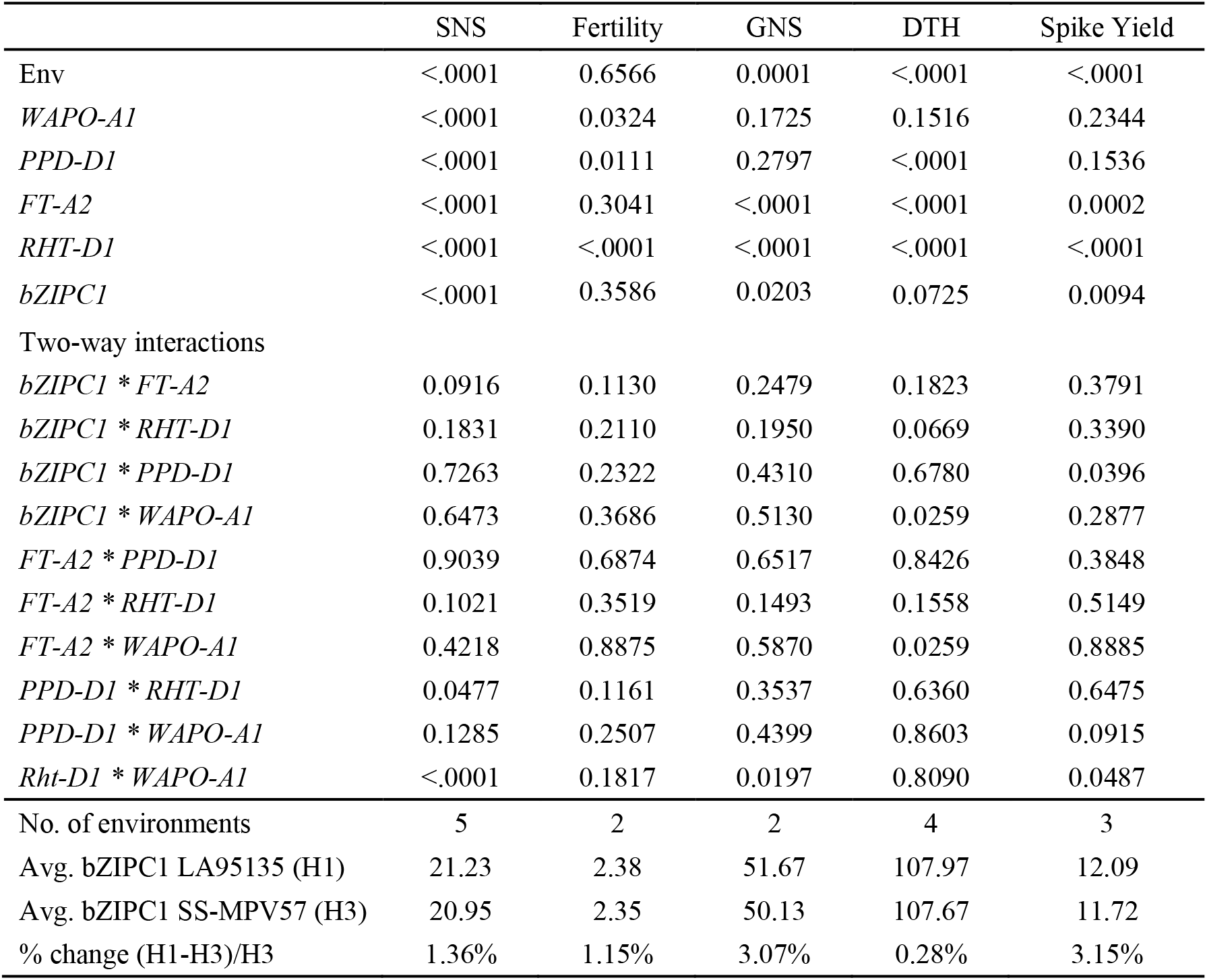
Factorial ANOVA for spikelet number per spike (SNS), grain number per spike (GNS), fertility, days to heading (DTH), and spike grain yield including 5 genes (factors), two alleles (levels), and environments (blocks). Raw data is presented in Table S11.

We also explored the effect of the H1 haplotype relative to H3 in the winter wheat population LA95135 (H1) x SS-MPV57 (H3) evaluated in five different environments in North Carolina, USA (DeWitt et al. 2021). In addition to *bZIPC-B1*, this population segregates for another four genes affecting SNS including *PPD-D1*, *RHT-D1*, *FT-A2* (Glenn et al. 2022), and *WAPO-A1* (Kuzay et al. 2019). We inferred the *bZIPC-B1* genotype from the consensus of two SNPs flanking this gene at 616,367,516 and 616,955,782, and used this information together with the genotypes of the other four genes in a factorial ANOVA including the five genes as factors, all gene interactions and environments as blocks (Table 5, raw data in Table S11).

The combined ANOVA across the five environments explained 44% of the variation in SNS and revealed highly significant differences in SNS for all five genes including *bZIPC-B1* (*P* < 0.0001, Table 5). *PPD-D1* and *WAPO-A1* showed the strongest effects, whereas *bZIPC-B1* had the weakest effect, with the Berkut allele showing on average a 1.4% increase in SNS (Table 5). The differences in SNS for *bZIPC-B1* were significant in three out of the five environments when the ANOVAs were performed separately by environment. None of the two-way interactions involving *bZIPC-B1* with the other four genes were significant, but the interaction between *bZIPC-B1* and *FT2* was the closest to significance (*P* = 0.0916, Table 5). When the same population was analyzed only for *bZIPC-B1* and *FT2*, the interaction was highly significant (*P =* 0.0004, Fig. S5a-b), with the *FT-A2* A10 – *bZIPC-B1* H1 allele combination showing the highest SNS.

We also analyzed the effect of *bZIPC-B1* on days to heading (DTH), fertility, GNS, and GWS in the same LA95135 x SS-MPV57 population. We found non-significant differences for DTH or fertility, but we observed significant increases of 3.07 % in GNS (*P* = 0.0203) and 3.15 % in GWS (*P* = 0.0094) associated with the H1 haplotype (LA95135) relative to the H3 haplotype (SS-MPV57, Table 5). The highest GWS was observed among the RILs combining the *bZIPC-B1* H1 and *FT-A2* A10 alleles (Fig. S5c).

Taken together, these results indicate that the derived *bZIPC-B1* KM allele (H1 haplotype) is associated with beneficial effects on SNS, GNS, and GWS relative to the ancestral NV allele, providing a possible explanation for the rapid frequency increase of the derived KM allele during wheat domestication and improvement (Table 3).

## DISCUSSION

### bZIP transcription factors are a large and functionally diversified family

The basic leucine zipper (bZIP) transcription factors (TF) are a large and diverse family present in eukaryotes from *Saccharomyces cerevisiae* to *Homo sapiens* (Deppmann et al. 2006). They are characterized by two conserved domains. The first one is a DNA-binding domain rich in basic amino acids, which is followed by a leucine zipper consisting of several heptad repeats of hydrophobic amino acids. This last region favors the formation of homodimers or heterodimers, resulting in multiple combinations with unique effects on transcriptional regulation (Deppmann et al. 2006). In plants, the bZIP transcription factors have a variety of roles ranging from hormonal responses, light signaling, nitrogen/carbon metabolism, biotic and abiotic stress, flower induction, development, and seed storage and maturation (Agarwal et al. 2019; Alonso et al. 2009; Alves et al. 2013; Baena-González et al. 2007; Choi et al. 2000; Fujita et al. 2005; Frank et al. 2018; Jakoby et al. 2002).

The plant bZIP transcription factors have diversified into 13 subfamilies (A-L and S) (Guedes Corrêa et al. 2008). The bZIP proteins from the C-group, which is the focus of this study, preferentially form heterodimers with other members of the C-group as well as with members of the S1-group (Ehlert et al. 2006). Known as the “C/S1 network”, this network has been shown to be involved in amino acid metabolism, stress response, and energy homeostasis (Alonso et al. 2009; Dröge-Laser and Weiste, 2018; Feng et al. 2021). The C-group bZIP genes in Arabidopsis (*bZIP9*, *bZIP10*, *bZIP25* and *bZIP63*) participate in a wide range of functions. For example, *bZIP63* has been shown to regulate the starvation response (Mair et al. 2015) and to play an important role in fine-tuning of ABA-mediated, sugar-dependent abiotic stress responses (Matiolli et al. 2011); whereas *bZIP10* was found to modulate basal defense and cell death (Kaminaka et al. 2006) and to work with glutathione to induce various heat shock proteins (Kumar and Chattopadhyay, 2018).

### Evolutionary and functional characterization of *bZIPC* genes in grasses

In grasses, the bZIP transcription factors of the C-group form a family of four paralogous clades, with groups bZIPC1 and bZIPC2 more closely related to each other than to the bZIPC3 and bZIPC4 groups (Fig. 1). The bZIPC1 clade includes previously studied proteins such as wheat SPA HETERODIMERIZING PROTEIN (TaSHP) (Boudet et al. 2019), barley BLZ1 (Vicente-Carbajosa et al. 1998) and maize OHP1 (Pysh et al. 1993), whereas the *bZIPC2* clade includes the wheat STORAGE PROTEIN ACTIVATOR (TaSPA= bZIPC2) (Albani et al. 1997), barley BLZ2 (Oñate et al. 1999), and maize OPAQUE2 (Pysh et al. 1993). The bZIPC3 and bZIPC4 subgroups have been less characterized.

TaSPA (= bZIPC2) was shown to be involved in the regulation of grain storage proteins (Albani et al. 1997) and in starch accumulation (Guo et al. 2020). This protein is orthologous to barley BLZ2, which interacts with BLZ1 (= bZIPC1) (Oñate et al. 1999). Both *BLZ1* and *BLZ2* are involved in the regulation of seed storage proteins (Vicente-Carbajosa et al. 1998). The orthologs of *bZIPC2* in maize *OPAQUE2* (*O2*) (Pysh et al. 1993) and rice *RISBZ1* (*OsbZIP58*) (Kawakatsu et al. 2009) were initially identified as positive regulators of grain storage protein. Similarly, the orthologs of *bZIPC1* in barley (BLZ1) and maize (OHP1 and OHP1b) have been shown to bind to the promoters of the genes encoding storage proteins promoting their activation (Vicente-Carbajosa et al. 1998; Zhang et al. 2015). *TaSHP*, however, has been suggested to act as a repressor of glutenin storage protein synthesis in wheat (Boudet et al. 2019). These results suggest that bZIP genes from the C-group can act both as transcriptional activators or repressors of storage proteins.

In addition to their role in the regulation of grass storage proteins, *bZIPC1* and *bZIPC2* genes have been shown to have additional functions. The rice ortholog of *bZIPC1*, *OsbZIP33* (Os03g58250), is involved in the resistance of abiotic stresses via ABA-dependent stress signal transduction, with transgenic rice plants overexpressing *OsbZIP33* showing significantly higher survival rates than their wildtype counterparts after dehydration treatment (Chen et al. 2015). The significant reduction in SNS detected in the *bzipc1* wheat mutants in this study demonstrates a previously unreported role of *bZIPC1* in the regulation of spike development.

The grass *bZIPC3* and *bZIPC4* genes, which are more distantly related to *bZIPC1* and *bZIPC2* (Fig. 1), have been less characterized. The three wheat *bZIPC3* homoeologs have been shown to be upregulated by cold (Tian et al. 2022), whereas *bZIPC-A4* (TraesCS6A02G154600 = Traes_6AS_F1CEB89EE.2) was upregulated after drought and heat treatments in wheat seedlings (Liu et al. 2015). Given the high level of expression of *bZIPC3* and *bZIPC4* genes in the wheat developing spike (Fig. 4), it would be interesting to determine if their mutants also show variation in spike-related traits.

In summary, the previous studies suggest some degree of sub-functionalization within the bZIP C-group clade. This sub-functionalization is reflected in their different expression profiles (Fig. 4) and in our Y2H results (Table 2), which showed that each of the four wheat bZIPC proteins can interact with different sets of proteins encoded by *FT-*like and *CEN-*like genes.

### bZIPC1 interacts with FT2 and regulates SNS

Our Y2H experiments showed that the FT2 and bZIPC1 proteins can physically interact with each other in yeast. Two indirect sources of evidence suggest that this interaction is biologically relevant. First, *bZIPC1* and *FT-A2* are both expressed in the developing spike at the same stages and are co-localized within the same cells in the IM (Figs. 4B-C). In addition, we observed a genetic interaction between the *bZIPC-B1* and *FT-A2* natural variants on SNS (Fig. S5), which may be related to the observed physical interaction between the proteins encoded by these two genes.

In spite of its sequence similarity with *FT1*, *FT2* shows some unique characteristics that set it apart from other *FT-*like genes. *FT2* is the only *FT*-like gene in wheat that is expressed directly in the developing spike (Shaw et al. 2019; VanGessel et al. 2022). It is also the only one that encodes a protein that does not interact with any of the known bZIP proteins of the A-group (FDL2, FDL6, and FDL15) or the known 14-3-3 proteins that interact with the other FT-like proteins (Li et al. 2015).

These unique characteristics motivated us to look for FT2 protein interactors and led to the discovery of bZIPC1 as an FT2 interactor. We initially hypothesized that knock-out mutants in *bZIPC1* would limit the ability of *FT2* to accelerate the formation of the terminal spikelet and would result in an increase in SNS as in the *ft2* mutants. However, all three *bzipc1* mutants in tetraploid wheat showed a significant decrease in SNS (Fig. 5), negating our initial hypothesis. We discuss below three alternative hypotheses that may explain the reduced SNS in the *bzipc1* mutant.

Based on the observed physical and genetic interactions between FT2 and bZIPC1 our favored hypothesis is that bZIPC1 interaction with FT2 may result in a non-functional protein complex that competes with other FT2 interactors required for FT2 function. Under this scenario, the *bzipc1* mutant could reduce competition with an interactor required for FT2 function, resulting in increased activity and reduced SNS. However, since bZIPC1 can also interact with FT3 and weakly with FT5 (Fig. 2), we cannot rule out alternative hypotheses in which the *bzipc1* effect on SNS is mediated by FT3 and/or FT5. Finally, *bZIPC1* may impact SNS by its effects on stress or nutrition pathways. This hypothesis is based on the known role of Arabidopsis *bZIP* genes from the C-group on the regulation of energy metabolism (Frank et al. 2018; Matiolli et al. 2011). Under starvation, the C/S_1_ network (Ehlert et al. 2006) reprograms metabolic gene expression to support survival by controlling plant growth, development and stress responses through fine tuning carbon and nitrogen responses (Dröge-Laser and Weiste, 2018). Since wheat plants affected by abiotic stresses or reduced nitrogen show reduced SNS (Frank and Bauer 1982; Frank et al. 1987; Maas and Grieve 1990), it will be interesting to investigate if wheat *bZIPC* genes are involved in these responses.

### Natural variation in bZIPC1 and potential applications to plant breeding

Analysis of the natural variation in the *bZIPC-B1* chromosome region revealed three major haplotypes (H1-H3), with the H1 haplotype associated with increases in SNS or GNS in four wheat RIL populations evaluated in multiple environments (Tables 4 and 5). The frequency of the H1 haplotype was higher in the durum landraces (37%) than in the ancestral domesticated Emmer (4%), and increased again in cultivated durum (65%) and common wheat (71%, Table 3). The historical increase in the H1 haplotype frequency and its favorable effect on SNS suggest that this haplotype has been favored by positive selection in wheat breeding programs. The current frequencies also indicate that there is still a significant number of common and durum wheat varieties that can potentially benefit from the incorporation of the H1 haplotype.

The bZIPC-B1 protein in the H1 haplotype differs from other haplotypes by two missense SNPs that result in two linked amino acid changes (KM in H1 and NV in the other haplotypes). Since loss-of-function mutations in this gene are associated with variation in SNS, we hypothesize that these amino acid changes can be the cause of the observed natural differences in SNS. However, a more precise genetic map of this trait will be required to rule out the possibility that other genes linked to *bZIPC-B1* contributed to the differences in SNS and the changes in allele frequency.

The two amino acid changes in the bZIPC-B1 protein may also affect its interactions with other proteins, and may explain the significant genetic interaction for SNS detected between *bZIPC-B1* and *FT-A2* in the RIL population LA95135 x SS-MPV57 (Fig. S5). In this population, the effect of *bZIPC-B1* on SNS was significant in the presence of the *FT-A2* A10 allele but not in the presence of the D10 allele. Similarly, the effect of *FT2* on SNS was significant in the presence of the *bZIPC-B1* H1 haplotype but not in the presence of the NV allele (Fig. S5). This epistatic interaction may be associated with the simultaneous and rapid increase of the *FT2* A10 (60-80%, Glenn et al. 2022) and *bZIPC-B1* H1 haplotype frequencies in modern common wheat cultivars, and suggests that simultaneous selection for these two alleles may be a useful breeding strategy to increase SNS.

In addition to its favorable effect on SNS, the H1 haplotype showed significantly higher GNS and GWS than the H2 and H3 haplotypes in multiple locations and in both spring and winter wheat. This is an encouraging result because it suggests that increases in grain number were not offset by a decrease in kernel weight. The interaction between *bZIPC-B1* and *FT-A2* for GWS was not significant but the trend was the same as for SNS, with the largest increase in spike weight detected in the RILs combining the *FT-A2* A10 and *bZIPC-B1* H1 alleles (Fig. S5c). The magnitude of the *bZIPC-B1* H1 haplotype effect on GWS and its interactions with *FT-A2* varied across environments, similar to what has been reported previously for other wheat genes affecting SNS such as *FT-A2* (Glenn et al. 2022) and *WAPO-A1* (Kuzay et al. 2019, 2022). Therefore, before deployment of these alleles into breeding programs, it would be prudent to test their effects in local genetic backgrounds and environments.

Fortunately, the favorable H1 haplotype in *bZIPC-B1* and the A10 allele in *FT-A2* are frequent in the wheat germplasm, which together with the haplotype analysis presented here, will facilitate the evaluation of the effect of these two alleles in breeding programs that have access to genotypic information. The diagnostic molecular markers developed in this study for the beneficial *bZIPC-B1* allele together with the markers previously developed for the *FT-A2* alleles (Glenn et al. 2022) are useful tools to evaluate the distribution and the effect of these alleles in a breeding program, and to accelerate their deployment. The sequences of these markers are publicly available to facilitate the deployment of these potentially beneficial alleles.

## STATEMENTS AND DECLARATIONS

### AUTHORS CONTRIBUTIONS

PG collected and analyzed most of the experimental data. DPW performed the Y2H screen, identified the bZIPC1 interactor, contributed to data analyses and supervised PG and NO. NO contributed to the Y2H screen, JZ contributed natural variation data and statistical analyses. GG provided valuable intellectual and writing contributions. JD and DPW initiated and coordinated the project. JD obtained funding and administered the project, contributed data analyses, supervised JZ, PG and DPW, and was responsible for the final manuscript. All authors reviewed the manuscript and provided suggestions.

### DATA AVAILABILITY

All data generated or analyzed during this study are included in the manuscript and supporting files. Kronos mutants are available from the authors upon request without any restrictions for use and from the Germplasm Resources Unit (GRU) at the John Innes Centre.

### ETHICS APPROVALS and CONSENTS TO PARTICIPATE

Do not apply to this study

## SUPPLEMENTAL DATA

### Supplemental Figures

**Figure S1.** Amino acid sequence alignment of bZIP C-group proteins.

**Figure S2**. Effects of splicing mutants K2991 and K3532 on *bZIPC-B1* transcripts.

**Figure S3**. Transcripts per million of the four *bZIPC* genes from the C-group.

**Figure S4**. Characterization of F_3_ homozygous mutants *bzipc1-1* and *bzipc1-2* (mut.) and their corresponding wildtype sister lines (WT).

**Figure S5.** *bZIPC*-*B1* x *FT-A2* interactions for spikelet number per spike and grain weight per spike in the LA95135 x SS-MPV57 RIL population.

### Supplemental Tables

**Table S1.** List of markers and primers used throughout the study.

**Table S2.** List of interactors identified in the yeast two hybrid (Y2H) screens.

**Table S3.** Effect of *bzipc1* mutants on SNS, heading time and, and leaf number (Fig. 5, F_2_).

**Table S4.** Effect of *bzipc1* mutants on SNS, heading time and, and leaf number (Fig. S4, F_3_).

**Table S5**. Haplotype bZIPC1 region.

**Table S6.** Distribution of *bZIPC-B1* non-synonymous SNPs among grasses.

**Table S7.** Frequencies of *bZIPC-B1* alleles in different wheat species and subspecies.

**Table S8**. Berkut (H1) x Patwin-515HP (H3).

**Table S9**. Berkut (H1) x RSI5 (H3).

**Table S10**. Berkut (H1) x Dharwar-Dry (H2).

**Table S11**. Winter wheat population LA95135 (H1) x SS-MPV57 (H3)

