## Supplemental Figures for "Wheat bZIPC1 interacts with FT2 and contributes to the regulation of spikelet number per spike"

**Figure S1.** Amino acid sequence alignments of bZIP C-group proteins. (a) Conserved basic region and leucine zipper (83 amino acids) used for the phylogenetic analysis in Figure 1. (b) c1 domain conserved between bZIPC1 proteins in grasses and Arabidopsis bZIP63.

(a)

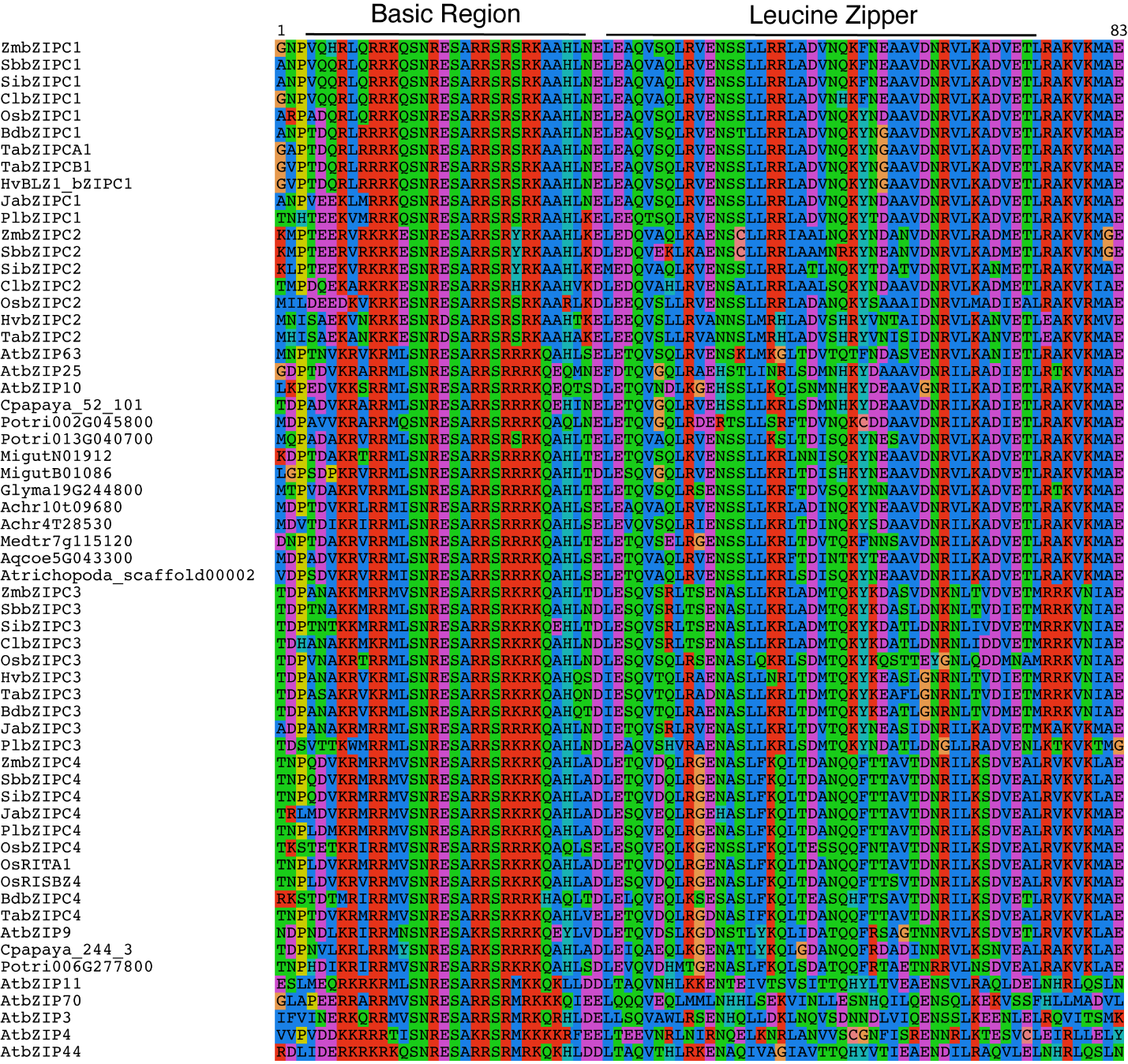

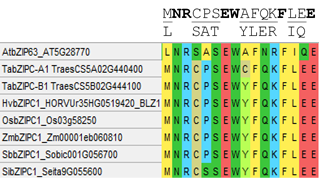

(b)

**Figure S2.** Effects of splicing mutants K2991 and K3532 on *bZIPC-B1* transcripts (primers in Table S1).

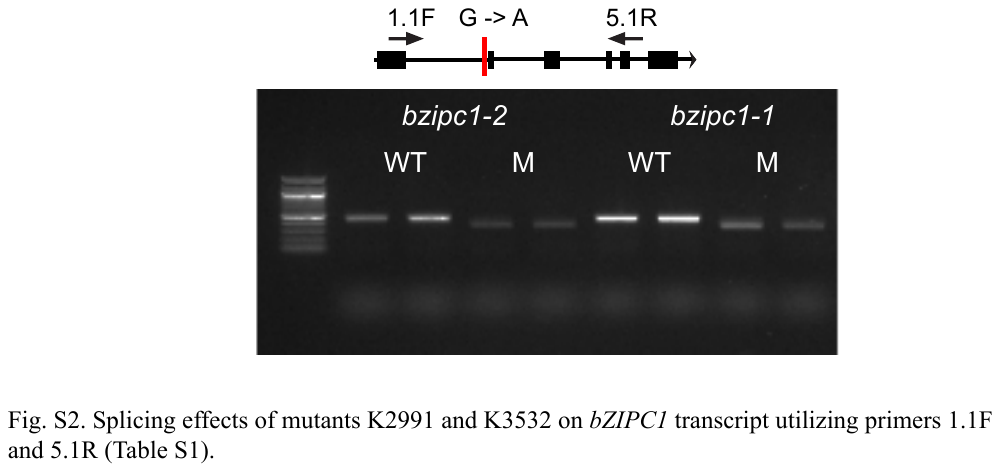

**Figure S3**. Transcripts per million (TPM) of the four *bZIPC* genes from the C-group from previously published Chinese Spring RNAseq expression across five tissues and three developmental stages (Choulet et al. 2014).

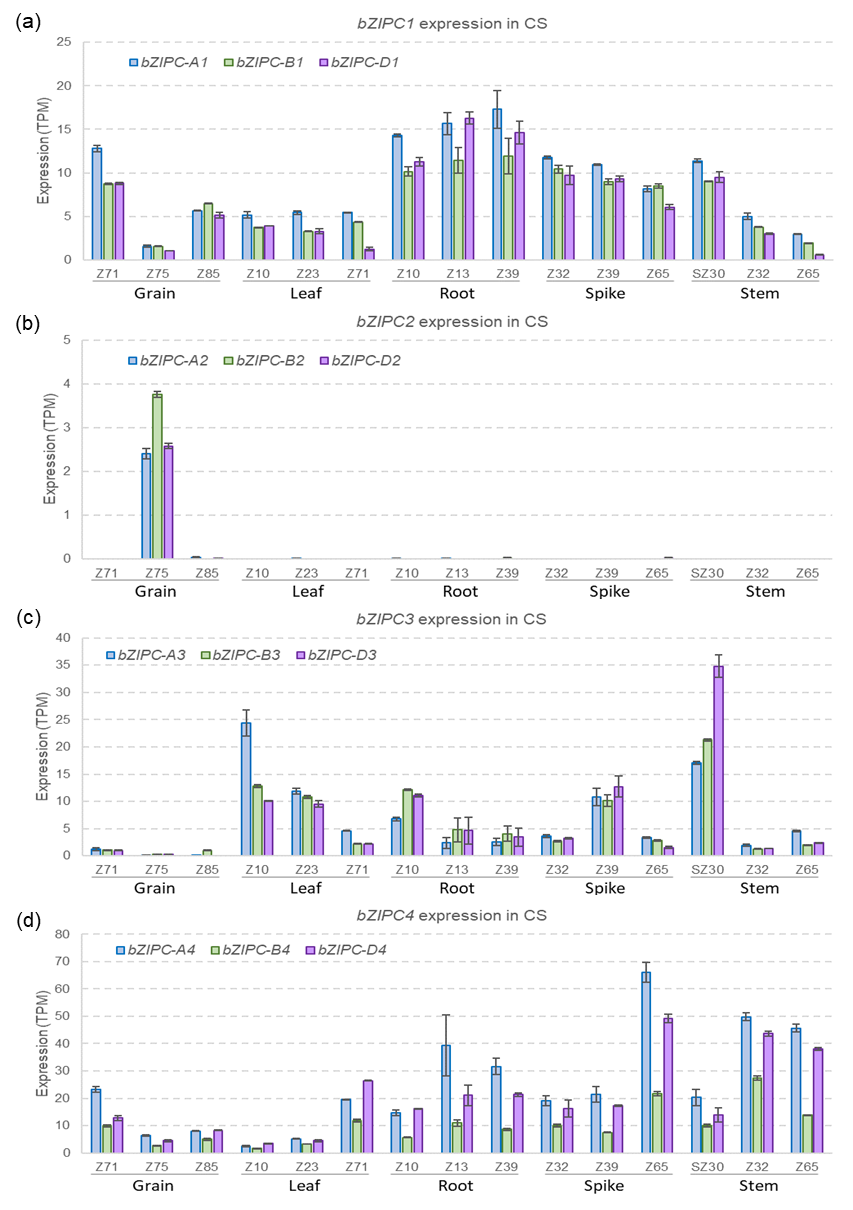

**Figure S4.** Characterization of F_3_ homozygous mutants bzipc1-1 and bzipc1-2 (mut) and their corresponding wildtype sister lines (WT). (a) Spikelet number per spike (SNS). (b) Days to heading (DTH). (c) Leaf number at heading (LN). ns= not significant, * = P < 0.05, ** = P < 0.01, *** = P < 0.001 based on two-tailed *t-*tests. N= 6 per genotype. Error bars are s.e.m. Raw data and statistical analysis are available in Table S4.

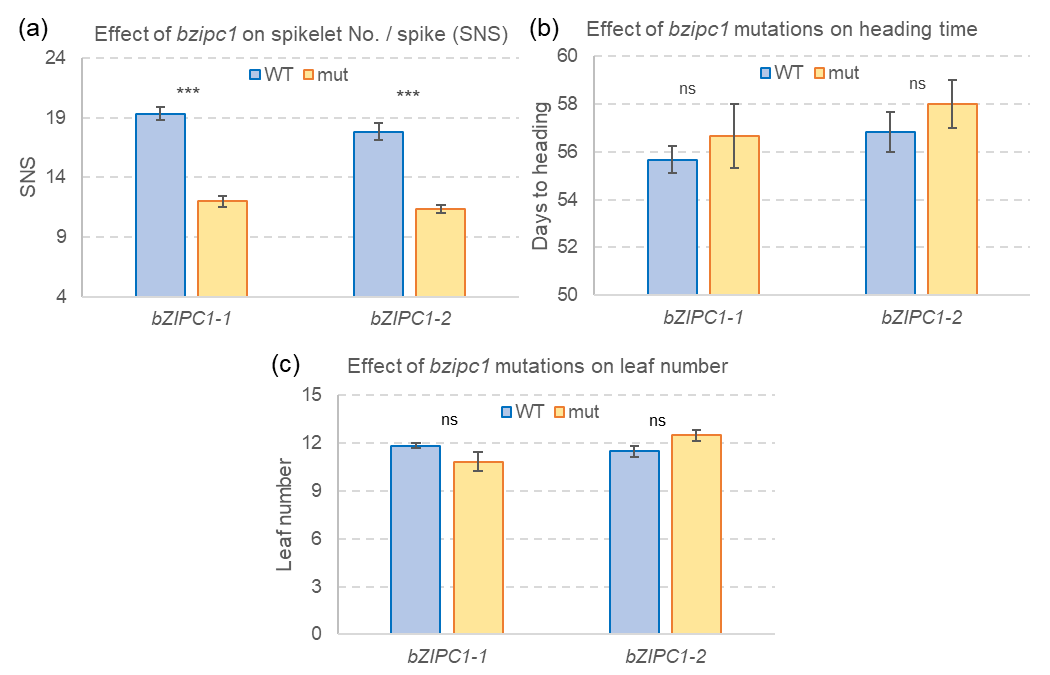

**Figure S5.** *bZIPC*-*B1* x *FT-A2* interaction for spikelet number per spike (SNS) and grain weight per spike (GWS) in the LA95135 x SS-MPV57 RIL population. (a) Interaction graph for the LSMEANS obtained in a factorial ANOVA including only *FT2*, *bZIPC1* and environment as factors. The four simple effects were performed using one-way ANOVAs by gene. Bars are standard errors of the least square (LS) means. ns= not significant, * = *P* < 0.05, *** = *P* < 0.001. (b) *P* values for the *bZIPC*-*B1* x *FT-A2* ANOVAs by environment. Note that in the factorial including only bZIPC1 and FT2 more data can be used (1,603 RIL/env.) than in the factorial including all genes (1,188 RIL/env.) due to the elimination of RILs with missing data in any of the genes for the factorial analyses. (c) Similar analysis for grain weight per spike. This trait is more variable than SNS and was evaluated in only three environments, with only one of them with close to significant differences (*P =* 0.0581). Although the overall interaction across environments was not significant (*P* = 0.2176), the trends were the same as for SNS: *i*) the *bZIPC1* H1 – *FT-A2* combination showed the strongest effect, *ii*) significant differences between *bZIPC-B1* were detected only within the *FT-A2* A10 allele, and *iii*) significant differences between the *FT2* alleles were detected only within the *bZIPC-B1* H1 allele. Raw data is available in Table S11.

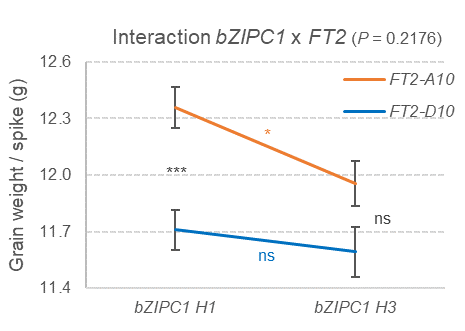

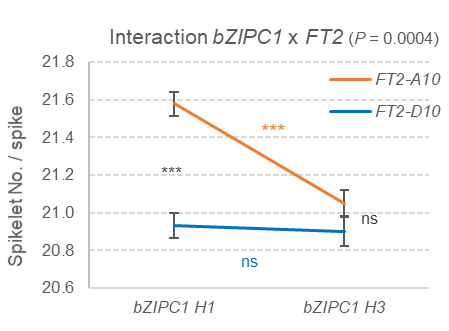

(a)

(c)

Interaction bZIPC1 x FT2 (P = 0.2176)

Grain weight / spike (g)

|  | FT2-A10 | FT2-D10 |
| --- | --- | --- |
| bZIPC1 H1 | 12.3598 | 11.7089 |
| bZIPC1 H3 | 11.9554 | 11.5931 |

Interaction bZIPC1 x FT2 (P = 0.0004)

Spikelet No. / spike

|  | FT2-A10 | FT2-D10 |
| --- | --- | --- |
| bZIPC1 H1 | 21.5773 | 20.9326 |
| bZIPC1 H3 | 21.0511 | 20.901 |

(b)

| ANOVAs for SNS including only *bZIPC-B1*, *FT-A2* and environment | | | | |
| --- | --- | --- | --- | --- |
| Environment | N used | FT-A2 | bZIPC-B1 | Interaction |
| Kinston, NC 2018 | 321 | 0.6315 | 0.0107 | 0.9142 |
| Kinston, NC 2019 | 319 | 0.0049 | 0.0241 | 0.0241 |
| Plains, GA 2019 | 321 | <0.0001 | 0.0295 | 0.1170 |
| Raleigh, NC, 2018 | 321 | 0.3566 | 0.2168 | 0.0817 |
| Raleigh, NC, 2019 | 321 | 0.0046 | 0.5714 | 0.0071 |
| 5 environments | 1,603 | <0.0001 | <0.0001 | 0.0004 |
| ANOVAs including bZIPC-B1, FT-A2, PPD-D1, RHT-D1, WAPO-A1 & env. | | | | |
| 5 environments | 1,188 | <0.0001 | <0.0001 | 0.0916 |
